# On the origin of an insular hybrid butterfly lineage

**DOI:** 10.1101/2024.10.17.618839

**Authors:** Jesper Boman, Zachary J. Nolen, Niclas Backström

**Author notes:** Emails: Jesper Boman, Zachary J. Nolen, Niclas Backström.

## Abstract

A new species can form through hybridization between species. Hybrid speciation in animals has been intensely debated, partly because hard evidence for the process has been difficult to obtain. Recent access to whole-genome sequencing data have made it more feasible to detect hybrid lineages. Here we report the discovery of a European hybrid butterfly lineage, a finding that can be considered surprising given the intense and long-term study of European butterflies. The lineage we describe is mainly inhabiting an island in the Baltic Sea and was previously designated as a subspecies (*horkei*) of one of the parental species (*Aricia artaxerxes*). By analysing whole-genome resequencing data, we determine that *horkei* originated by hybridization between the non-sister species *A. artaxerxes* and *A. agestis*. This hybridization event occurred approximately 54,000 years ago, predating the last glaciation of the current distribution range. *Horkei* must therefore have persisted long enough to be able to colonize its current range, despite that this area lies between the current distributions of the parental species. The hybrid origin, the maintenance of genomic integrity across times of dramatic climate change and the expression of a combination of parental traits suggest that *horkei* could be in the process of hybrid speciation.

## 1. Introduction

Hybridization between species has traditionally been considered rare and of limited importance in animal evolution. However, recent studies of hybrid zones and analyses of whole-genome resequencing data have challenged this view. In several different animal systems, hybridization has been shown to mediate exchange of adaptive alleles [1–3]. Even so, the majority of alleles introgressed from a divergent lineage are expected to be deleterious in the recipient population [4,5]. A more profound outcome of hybridization is the formation of a hybrid lineage or species [6–8]. For a hybrid species to persist, it needs to be (at least partially) reproductively isolated from both parental species. This can be achieved if hybridization causes a change of ploidy, leading to partially sterile offspring when hybrids backcross with either of the parental species [9]. However, ploidy shifts are much rarer in animals than in plants and homoploid hybrid speciation is thus a more likely mechanism for the evolution of animal hybrid species [10]. Exactly how frequently homoploid hybrid speciation occurs is unknown for several reasons. Post-speciation gene-flow may for example erase most of the ancestry of one of the parental species and make a hybrid origin difficult or impossible to infer [11]. Consequently, it is easier to verify hybrid origin for more recently established lineages, where ancestry patterns are more straightforward to infer, enabling testing for hybrid origin. With whole-genome sequencing data we can now identify hybrid ancestry, systematically test for the likelihood of a hybrid origin and infer how long a hybrid lineage has persisted.

In butterflies, several potential cases of homoploid hybrid speciation have been described [11– 16]. Here, we study hybridization between the northern brown argus (*Aricia artaxerxes*) and the brown argus (*Aricia agestis*), two butterfly species belonging to the blues (Lycaenidae). The two species differ slightly in morphology, with *A. agestis* tending to show additional and more pronounced orange spots on the dorsal side of the wings, but intraspecific variation exists and this character is not entirely diagnostic (Figure 1A) [17–19]. However, the species also differ in voltinism, with *A. artaxerxes* having one generation per year (univoltine) with a peak occurrence in between the two generations of the bivoltine *A. agestis* (Figure S1). This is important, since the difference in voltinism may underlie the geographic distribution gaps that are sometimes observed between *A. artaxerxes* and *A. agestis* [20]. Northern European and British *Aricia* were the focus of several studies during the 20^th^ century [e.g. 17,21,22]. The main conclusion from that work was that *A. artaxerxes* and *A. agestis* should be considered different species, with the former having a more northern distribution and the latter being restricted to the southern regions of e.g. Sweden and Britain. Over the last decades, *A. agestis* has expanded northwards in Britain [23]. A genetic analysis of two loci revealed that *Aricia* in Northern England and Wales likely have hybrid ancestry, resulting from historical hybridization between *A. agestis* and *A. artaxerxes* [24]. In Sweden, both species are rather common, and they have almost disjunct distributions. *Aricia agestis* occurs in the extreme south but has recently expanded somewhat along the coastal areas in the south-east. *Aricia artaxerxes* is widely distributed throughout most of the country, except in the far south and south-west, where it is rare or absent. On Öland, an island in the Baltic Sea, a subspecies of *A. artaxerxes (A. a. horkei)* has been described. *Aricia artaxerxes horkei (*from here on ‘*horkei*’) has befuddled researchers by having *A. agestis-*like morphology and host plant preference, but being univoltine like *A. artaxerxes* and interfertile in crosses with mainland *A. artaxerxes* [22]. In a genetic analysis of ten allozyme loci, *horkei* was placed as a sister clade to populations of *A. artaxerxes* from Britain and northern Scandinavia [19]. The conflicting data led us to hypothesize that *horkei* is a population with hybrid ancestry, like the *Aricia* in Northern England. We thus collected both *A. artaxerxes* and *A. agestis* in their core range in mainland Sweden, as well as *horkei* on Öland, with the aim to apply genomic methods to investigate the ancestry of *horkei*. In addition, we wanted to assess if the secondary contact zone formed by the range expansion of *A. agestis* in southeastern Sweden has resulted in gene flow between the two species.

**Figure 1.**
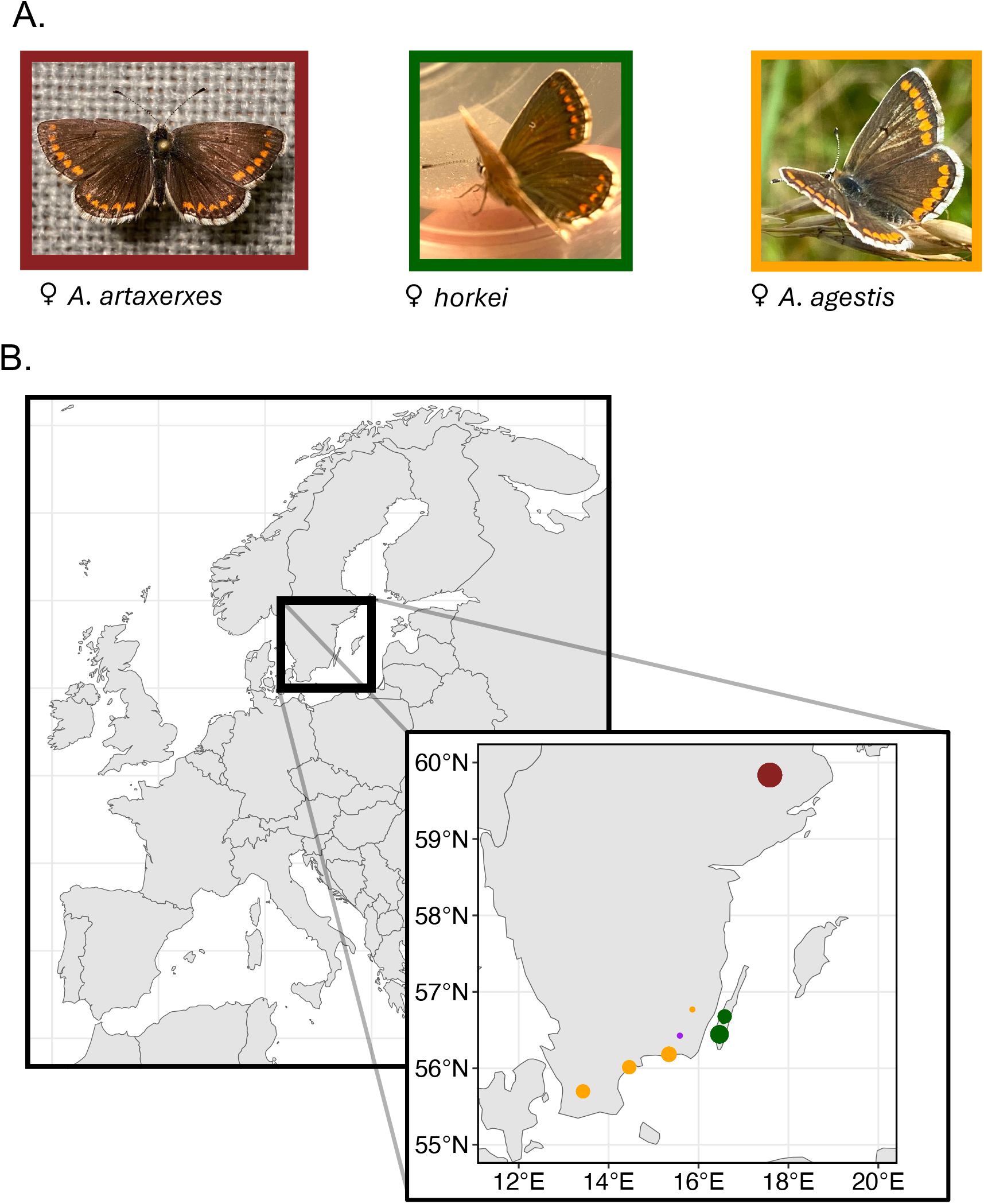
Photographs of representative females from the three taxa and a map of sampling locations. **(A)** Female individuals of *A. artaxerxes, horkei* and *A. agestis*, illustrating that *A. agestis* individuals in general have a higher number of, and more pronounced, orange dots (lunules) on the dorsal sides of both forewings and hindwings than *A. artaxerxes. Horkei* have on average an intermediate number of orange lunules [19]. **(B)** Map of Southern Sweden with sampling locations (coloured points). Points are scaled according to the number of samples collected at each site, ranging from n = 1 to n = 17. Colour code in **(B)** follows background colour of photographs in **(A)**. In addition, a *horkei* individual from the mainland (“Blekingehorkei”) is shown in purple in (**B**).

## 2. Results

### Sampling and sequencing

We sampled *Aricia* individuals during the summer of 2022. Southeastern Sweden (including Öland) was visited late in the flight period for *A. artaxerxes* and early in the second-generation flight period of *A. agestis* to include the possibility of co-occurrence (Figure 1B and Table S1). We sampled one *A. agestis* individual in Nybro, Småland which is the northernmost genetically confirmed record of this species in Sweden, supporting the view that this species is expanding north. We did not find any populations in sympatry but discovered a mosaic distribution with the closest localities of each species separated by 30 kilometres. In total, we whole-genome sequenced 45 individuals using Illumina short reads from both species, including 13 *horkei* from Öland, as well as 17 *A. artaxerxes* from central Sweden, one from the south-east and 14 *A. agestis* in the south and south-east (Figure 1B). Mean read depth across samples was 30.4x.

### Population genetic structure analyses

We used 3,378,264 high-quality autosomal single-nucleotide polymorphisms (SNPs) to investigate population structure. The principal component analysis showed that the first axis explained 22 % of the variation and differentiated *A. artaxerxes* from central Sweden and *A. agestis*, with an intermediate placement of *horkei* (Figure 2A). The second principal component mainly separated *horkei* and one individual collected in Blekinge in the south-east (hereafter Blekingehorkei, Figure 1B) from all other individuals. As an additional method we performed ADMIXTURE analyses (Figure 2B), assuming K=1-10 clusters of ancestry. The cross-validation error showed a monotonic increase from K=1 to K=10, highlighting that much genetic variation is shared across species (Figure S2). Assuming K=2, a similar pattern to the principal component analysis emerged with *horkei* having 68-70 % *artaxerxes* ancestry and Blekingehorkei 77 % (Figure 2B). Using K=3, *horkei* is differentiated as a separate cluster with Blekingehorkei as an intermediate between *horkei* and *A. artaxerxes* (Figure 2B). These results suggests that *horkei* has a mixed ancestry from both parental species and thus constitutes a hybrid lineage. The results also indicate that Blekingehorkei stems from a secondary admixture event with *artaxerxes*.

**Figure 2.**
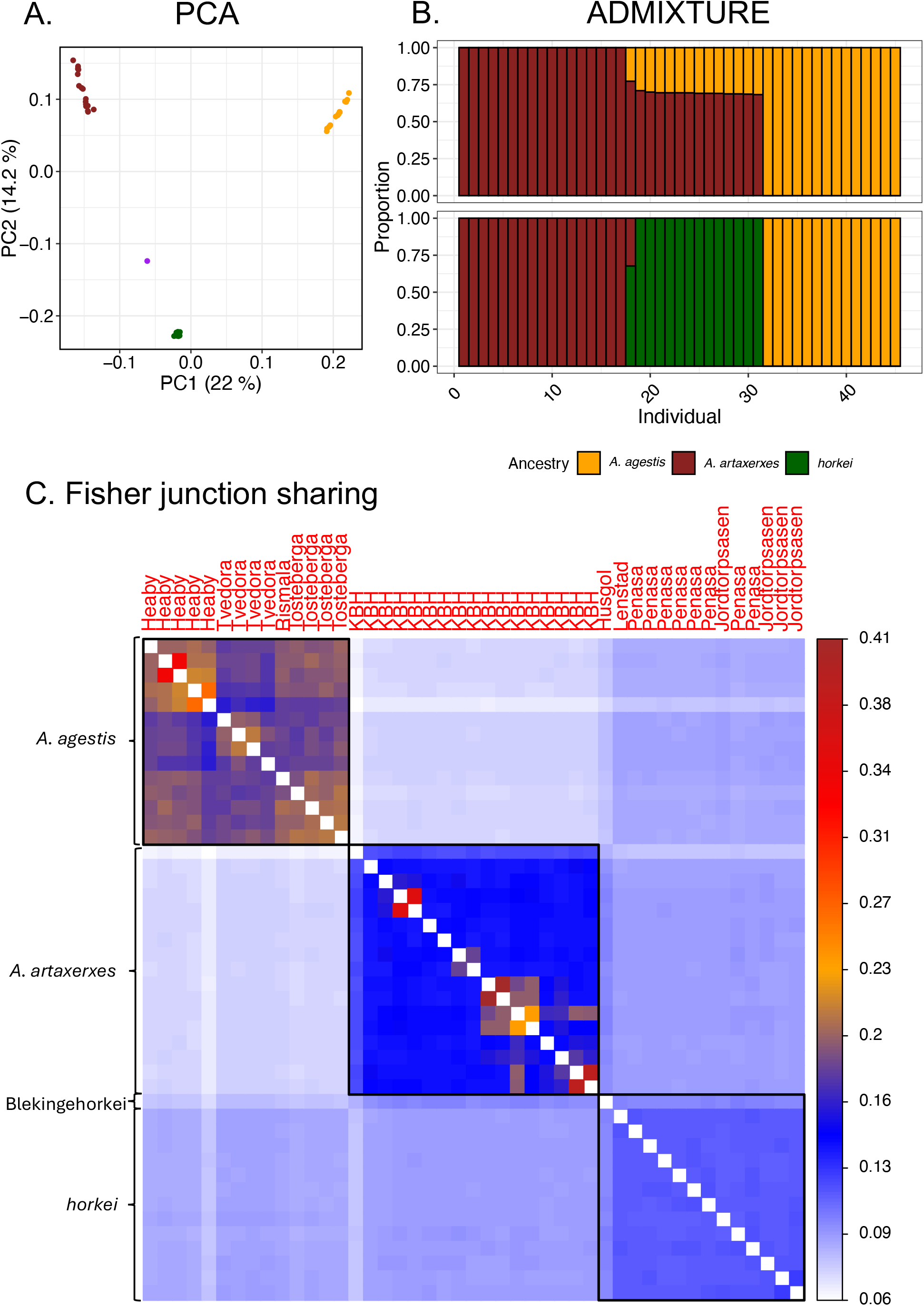
Population genetic cluster analyses. (**A)** The results from the principal component analysis of high-quality SNPs. **(B)** Results from the ADMIXTURE analysis for K=2 and K=3. K=1 had the lowest cross-validation score (Figure S2). **(C)** Genetic structure based on Fisher junction sharing. We developed a novel method, based on the 70-year old theory of Fisher junctions, to analyse population genetic structure. We calculated the sharing of Fisher junctions for all pairwise relationships between samples and used hierarchical clusters to order the samples in the plot. Major clusters are shown as black-bordered boxes. Samples predominantly group by species/population (Y-axis labels) and, within species, most samples cluster according to sampling locality (X-axis labels). Notably, *horkei* and Blekingehorkei share more Fisher junctions with each other than either of them does with *A. agestis* or *A. artaxerxes*. The legend below the ADMIXTURE plot illustrates the colour codes for each respective species (applies to panel **A** and **B**). Blekingehorkei is shown in purple in (**A**).

### Fine-scale population structure revealed by sharing of Fisher junctions

Patterns of ancestry across the genome of admixed individuals occur in so called haplotype blocks. Here, we define a haplotype block as a genomic segment that is identical-by-descent (IBD) in our set of genome sequences [see 25 for an alternative definition]. Distribution of ancestry-specific haplotype block lengths hold information e.g. on timing of hybridization. However, correctly inferring haplotype block lengths is difficult and can be error prone (haplotype switch errors) [26]. With some loss of information, we can instead use the so-called Fisher junctions that separate adjacent haplotype blocks to make population genetic inferences [27,28]. Two individuals are more closely related genetically if a larger fraction of the genome is IBD, by definition. Concordantly, two individuals are more closely related if they share a larger fraction of Fisher junctions. We applied this concept to our data by first phasing high-quality biallelic SNPs using SHAPEIT: a statistical phasing method that essentially reconstructs haplotype blocks based on patterns of linkage disequilibrium between loci (Figure S3). We then defined Fisher junctions as the boundaries between haplotype blocks (Figure S3), calculated all pairwise shared Fisher junctions and clustered individuals based on the pairwise relationships. In line with the results from ADMIXTURE, this analysis revealed three well-separated clusters, corresponding to *A. agestis, A. artaxerxes* and *horkei* (including Blekingehorkei; Figure 2C). In addition, the Fisher junction analysis showed that *A. agestis* individuals clustered by sampling locality and individuals within localities clustered by relatedness (Figure 2C and Figure S4). Little variation existed between *horkei* individuals in their degree of junction sharing with parental species. This is in line with a lack of recent gene flow between *horkei* and either of the parental species, as suggested from the ADMIXTURE analysis (Figure 2B). Blekingehorkei shared slightly more junctions with *horkei* (0.098) than with *A. artaxerxes* (0.092) or *A. agestis* (0.077). This suggests that the *A. agestis* ancestry in Blekingehorkei can be traced back to the same hybridization event that led to the origin of *horkei* and is in line with the results from the ADMIXTURE analysis. To further confirm this, we inferred fixed differences between *A. agestis* and *A. artaxerxes* (i.e. excluding *horkei* and Blekingehorkei) and found 5,759 autosomal fixed differences. Of these, Blekingehorkei only have 7 *A. agestis* alleles absent in the *horkei* population (Figure S5), in line with expectations for a common ancestry of *A. agestis*-alleles in Blekingehorkei and *horkei*.

### Island hybrids are derived from a single ancestral mitochondrial lineage

We next considered whole mitochondrial genomes (mtDNA) to gain further insights on the origin of *horkei*. A maximum likelihood phylogeny revealed that *A. agestis* and *A. artaxerxes* are reciprocally monophyletic, and that *horkei* is nested within artaxerxes (Figure 3). *A. artaxerxes* and *A. agestis* mitochondrial genomes differ at approximately 400 sites (i.e. net mtDNA divergence is ∼2.6%). Notably, *horkei* forms a monophyletic lineage within *A. artaxerxes*, suggesting that all *horkei* are derived from a single mitochondrial lineage. The mitochondrion of Blekingehorkei was most closely related to an *A. artaxerxes* individual sampled in central Sweden (Uppland, Figure 1B).

**Figure 3.**
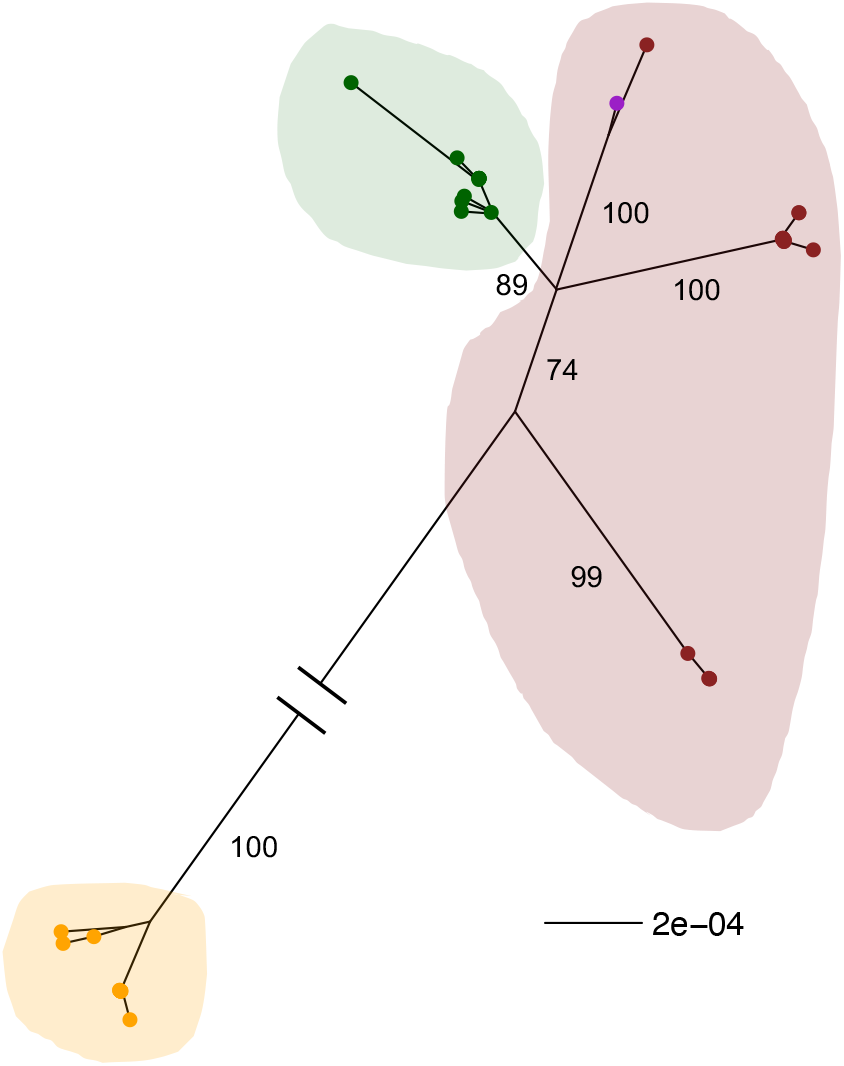
A maximum likelihood phylogeny based on whole (∼15.5 kb) mitochondrial genome sequences. The long branch (∼ 402 substitutions) between *A. agestis* (orange) and *A. artaxerxes* (brown) has been truncated for the purpose of presentation. *Horkei* samples are shown in green and Blekingehorkei in purple (nested within the *artaxerxes* group). The scale bar represents the number of substitutions per base pair.

### Multispecies coalescence analysis strongly supports hybrid origin of *horkei*

To infer the evolutionary history of *horkei*, we fit multispecies coalescence with introgression models to 1,000 autosomal loci using BPP [29]. All four models predicted that the split between *A. agestis* and *A. artaxerxes* occurred ∼415 k generations ago. The hybridization (or introgression) giving rise to the admixed *horkei* was inferred to have happened between 37 k (Model A) and 54 k (Model C) generations ago (Figure 4A and Table 1). The inferred ancestry proportions of each parental lineage in *horkei* ranged between 42-57% across models. Despite these consistent results, an analysis of marginal likelihoods showed that Model C fitted significantly better to the data than any other model (Table 1), supporting that *horkei* has a hybrid origin. The odds ratio favouring Model C over Model B_Arx_, which had the second highest marginal likelihood, was 5.9*10^47^. In conclusion, our analysis suggests that *horkei* formed as a hybrid between the non-sister species *A. agestis* and *A. artaxerxes* (Figure 4B).

**Table 1.** Parameter estimates for the hybrid lineage formation model (Model C in Figure 4A). *N* represent the effective population size estimated from θ-values obtained by the model. Here *N* = θ/4μ, where μ is the mutation rate per base per generation. Times (τ_Hybrid formation_ and τ_Root_) are given in generations. These were obtained by dividing the coalescent units (C) with the mutation rate: C/μ. Currently, *horkei* has one generation per year, suggesting that hybrid formation (τ_Hybrid formation_) occurred roughly 50 - 58 k years ago. HPD = highest posterior density interval.

| Parameter | Mean | 95 % HPD: lower | 95 % HPD: upper |
| --- | --- | --- | --- |
| $N_{\text{Artaxerxes}}$ | 572,069 | 528,103 | 614,052 |
| $N_{\text{Horkei}}$ | 1,032,931 | 951,207 | 1,114,397 |
| $N_{\text{Agestis}}$ | 201,724 | 186,810 | 216,552 |
| $N_{\text{Artaxerxes - ancestor}}$ | 2,215,000 | 2,070,259 | 2,357,759 |
| $N_{\text{Agestis - ancestor}}$ | 3,009,310 | 2,758,448 | 3,237,586 |
| $N_{\text{Common ancestor}}$ | 808,793 | 781,466 | 834,914 |
| $\tau_{\text{Hybrid formation}}$ | 54,482 | 50,344 | 58,275 |
| $\tau_{\text{Root}}$ | 414,482 | 403,793 | 425,172 |
| $\phi_{\text{Proportion artaxerxes}}$ | 48 % | 45.3 % | 50.8 % |

**Table 2.** Bayesian estimates of marginal log likelihoods for the four different models describing the origin of *horkei*. The likelihoods were estimated with the thermodynamic integration model, using 16 separate runs of each model and sampling from the prior to the posterior.

| Model | Marginal log likelihood |
| --- | --- |
| A | -4.28308*10<sup>6</sup> |
| B<sub>Age</sub> | -4.28332*10<sup>6</sup> |
| B<sub>Arx</sub> | -4.28252*10<sup>6</sup> |
| C | -4.28241*10<sup>6</sup> |

**Figure 4.**
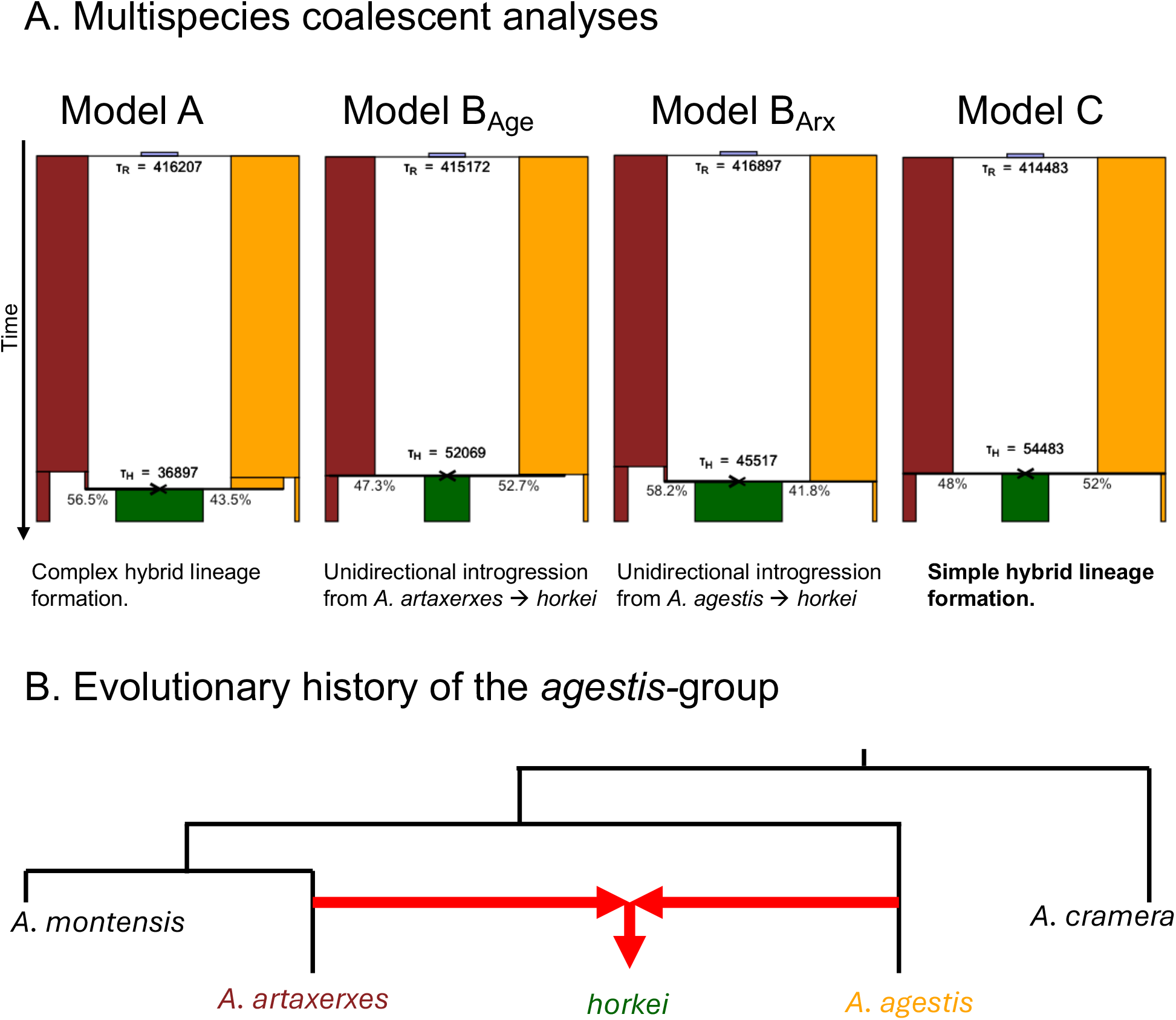
Multispecies coalescent models and cladogram of the *agestis*-group. **(A)** Schematics showing the multispecies coalescent models used to investigate the evolutionary history of *A. artaxerxes, A. agestis* and *horkei*. These are the four standard multispecies coalescence with introgression models suggested by BPP [29]. Model A represents a complex hybrid lineage formation, which can be interpreted either as introgression into the ancestor of *horkei* from an unsampled/ghost lineage branching from the other species, or hybrid species formation of two unsampled/ghost lineages. Model B_Age_ represents a scenario where *horkei* branched off from *A. agestis* and later received introgression from *A. artaxerxes*. Model B_Arx_ represents a scenario where *horkei* branched off from *A. artaxerxes* and later received introgression from *A. agestis*. Model C represents the simpler hybrid lineage formation scenario, in which *horkei* originated as a result of hybridization between *A. agestis* and *A. artaxerxes*. Boxes are scaled according to population size (N). The populations are from left to right for each model: *A. artaxerxes* (brown), *horkei* (green) and *A. agestis* (orange). The ancestral population (in the top middle) is shown in light blue. **(B)** Illustration of the evolutionary history of the *agestis-*group based on the multispecies coalescence analysis and a phylogeny from Sañudo-Restrepo et al., [30].

### Lack of *A. agestis* ancestry on the Z sex chromosome of *horkei*

When a hybrid population forms, parental haplotypes will segregate in the population. This will result in a higher level of genetic diversity than expected from the hybrid population size (*N*), if the parental lineages have some level of genetic divergence. With time, the hybrid population will reach an equilibrium genetic diversity which is determined by the strength of genetic drift (1/2*N*) and the mutation rate. To get a comparative estimate of the genome-wide level of genetic diversity, we calculated average pairwise differences (π) in 10 kb non-overlapping windows (Figure 5A). *Horkei* had a significantly higher π (0.0099, 95 % confidence interval [CI]: 0.0099 – 0.01) than both *A. artaxerxes* (0.0086, [0.0086 – 0.0086]) and *A. agestis* (0.0089, [0.0089 – 0.009]) (*p <* 0.05 in both cases). Visual inspection shows that the elevated π in *horkei* is obvious on all autosomes, but not on the Z chromosome (Figure 5A). To assess how the divergence process between *horkei* and the parental lineages has manifested in genome-wide patterns of genetic differentiation, we also estimated *F*_ST_ between *horkei* and the two parental populations in 10 kb non-overlapping windows. The level of genetic differentiation was significantly higher (*p <* 0.05) between *horkei* and *A. agestis* (mean: 0.1505, [0.1493 – 0.1516]) than between *horkei* and *A. artaxerxes* (0.0752, [0.0746 – 0.0757]). This pattern was especially pronounced on the Z chromosome, where *F*_ST_ was 0.4086 [0.4036 – 0.4135] between *horkei* and *A. agestis* and 0.1147 [0.1122 – 0.1171] between *horkei* and *A. artaxerxes*. Genetic differentiation between parental species was significantly (*p* < 0.05) higher on the Z sex chromosome (0.4593 [0.4548 – 0.4639]) compared to the genome-wide level (0.2125 [0.2112 – 0.2137]) (Figure S6).

**Figure 5.**
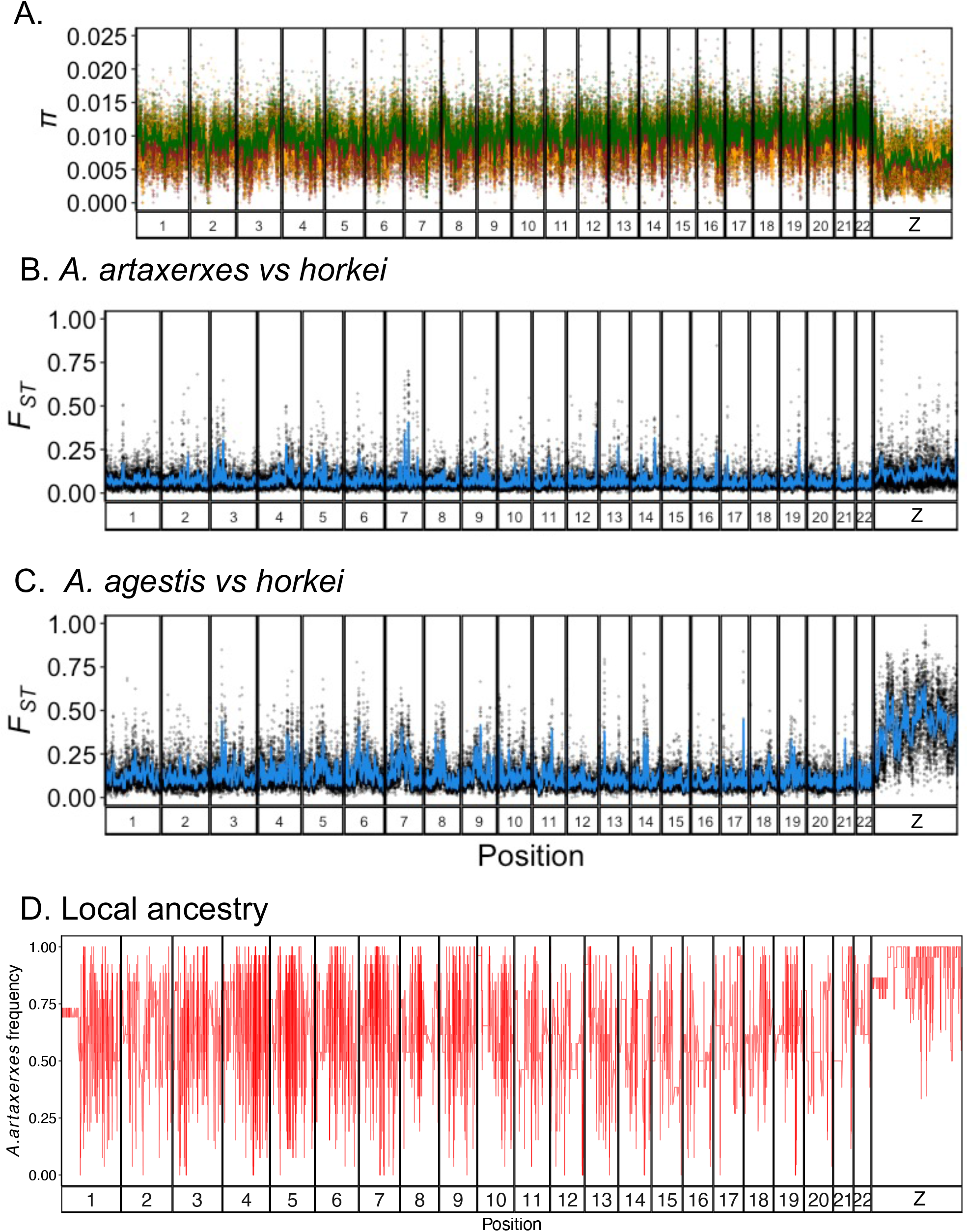
Genome-wide patterns of diversity, differentiation, and ancestry. **(A)** Genetic diversity (π) in 10 kb non-overlapping windows. Shown are *horkei* (green), *A. artaxerxes* (brown) and *A. agestis* (orange). **(B-C)** Genetic differentiation (*F*_ST_) between *horkei* and parental populations. Lines in **(A-C)** are local regression curves with the shaded region being the 95 % confidence interval. **(D)** *A. artaxerxes* local ancestry along the genome inferred using a hidden Markov model.

To gain further insights into the regional distribution of genetic ancestry proportions in *horkei*, we applied a method that infers local ancestry. Robust estimates of local ancestry require information about the variation in recombination rate. We therefore developed a genetic map for *A. artaxerxes* based on a pedigree consisting of a parental pair and 40 of their offspring. The inferred male-specific genetic map had a total length of 11.76 Morgans which translates to roughly one crossover per chromosome per meiosis (Figure S7). The recombination map was used to inform the hidden Markov model that was applied to infer the likelihood of ancestry switches along the genome [31]. The average frequency of *A. artaxerxes* ancestry for autosomes varied from 53% to 71% across *horkei* individuals (Figure 5D). For the Z chromosome, the average *A. artaxerxes* ancestry was significantly higher (on average 93%, Wilcoxon test *p <* 0.05). For the autosomes, 620 markers (1%) were fixed for the *A. agestis* allele and 2,016 (3.2%) for the *A. artaxerxes* allele. These proportions were skewed on the Z chromosome where no positions were fixed for the *A. agestis* allele while 2,352 markers (47%) were fixed for the *A. artaxerxes* allele. Taken together, this clearly shows that the autosomes in general show mosaic patterns of ancestry, while the Z sex chromosome is dominated by *A. artaxerxes* ancestry in *horkei*.

## 3. Discussion

### Population genetic structure of *Aricia* in southern Sweden

We used whole-genome resequencing data and evolutionary inferences to determine that *horkei* is not simply a subspecies of *A. artaxerxes*, but instead a stable hybrid lineage between the non-sister species *A. artaxerxes* and *A. agestis*. While it was previously thought that *horkei* was restricted to the island of Öland we here provided evidence that *A. artaxerxes* in the nearby province of Blekinge (Blekingehorkei) is more closely related to *horkei* than to *A. artaxerxes* from central Sweden. A reasonable interpretation is that Blekingehorkei has further admixed with *A. artaxerxes*, confirming incomplete reproductive isolation between *horkei* and mainland *A. artaxerxes-*like individuals from Skåne (ssp. *rambringi*) [22], which themselves may very well genetically be similar to Blekingehorkei. In the genome of the Blekingehorkei individual we only found 7 *A. agestis* alleles absent in the *horkei* samples. This small number can easily be due to sampling effects or loss of *A. agestis* alleles in *horkei* and we thus interpret this as lack of evidence for modern gene flow between *A. artaxerxes/horkei* and *A. agestis*. However, since we sampled Blekingehorkei and *A. agestis* individuals on consecutive days at localities 30 kilometres apart, possibility for hybridization may exist and will be interesting to follow up in future studies with a more comprehensive sampling. This is also noteworthy in light of the expansion of *A. agestis* into Småland, which has previously been documented along the coast (Cederberg, *personal communication*) and which we now confirm with an individual further inland. Interestingly, this *A. agestis* individual is more closely related to individuals from the province of Skåne than to the more geographically proximal individuals in Blekinge.

### Is *horkei* a distinct species?

Previous studies have established a close relatedness between *A. artaxerxes* and *A. agestis* [e.g. 17,19,21,22,24,30]. Our analysis suggests that they diverged around 400 Kya and that *horkei* originated around 54 Kya, predating the last time the Baltic Sea island of Öland was covered in ice during the last glacial maximum [32]. This indicates that the hybridization event leading to the formation of *horkei* did not occur on Öland. This is important since it is likely that the current allopatric/parapatric distribution of *horkei* with regards to the parental species does not reflect historical distributions. Thus, according to the model results, *horkei* must have reached Öland without the minor parent ancestry (*A. agestis*) being swamped by *A. artaxerxes* introgression. This gives us some information on how to interpret this enigmatic group of butterflies. It is possible that *horkei* and the Northern England *Aricia* arose from the same hybridization event. Both occupy latitudes that in modern times have been intermediate between *artaxerxes* and *agestis*. They both share the characteristic of a single brood per year (univoltine). On Öland this could be adaptive since the late spring on the island may not allow for two successful broods per year. Intriguingly, *horkei* on Öland most likely only uses rock roses (*Helianthemum spp*.) as host plants, like *A. agestis*, but unlike mainland *A. artaxerxes* further north that likely only use cranesbills (*Geranium spp*.) [33,34]. Previous observations indicate that *horkei* is largely absent from areas with *Geranium*, but common in areas with *Helianthemum*, and in localities where both plants are available, oviposition occurs exclusively on *Helianthemum* [22]. Thus, it appears that *horkei* has intermediate morphology (wing patterns, see *Introduction*) and ecological characteristics from both of the parental populations allowing it to occupy and thrive in the unique open alvar landscape on Öland [35,36]. With these facts in mind, we can now turn to the question of whether *horkei* is a species distinct from *A. agestis* and *A. artaxerxes*. According to a strict interpretation of the biological species concept which separates species based on reproductive isolation, *A. agestis* and *A. artaxerxes* should not be considered different species [18]. They are able to hybridize and produce viable offspring both in nature [this study and 24] and in the lab [17,21]. If we allow for species distinction based on partial isolation and focus on habitat isolation or ecogeographic isolation, i.e. differences in distribution driven by genetic differences [37], then all three main groups of this complex appear to be partially isolated with both *horkei* and Northern England *Aricia* occupying an intermediate niche. An attractive option here is to consider the genotypic cluster species concept, which emphasizes that species should maintain genetic integrity (at least in certain genomic regions) even if hybridization and gene flow with other species occurs [38]. Our results show that *horkei* can interbreed with *A. artaxerxes*, but the persistence as a distinct lineage since long before the colonization of Öland makes it reasonable to consider *horkei* a separate species: *Aricia horkei* [17]. Future evidence could either support or refute this status, which we discuss more below. We also want to emphasize that it might be possible that *horkei* and Northern England *Aricia* arose from the same hybridization event and could thus potentially be considered conspecific. We caution though that distinguishing a singular period of hybridization giving rise to both, may be difficult from two separate hybridizations occurring at similar timepoints from the same two large source populations. Nevertheless, there is some evidence that Northern England *Aricia* and *horkei* may be distinct, since the former has predominantly *A. agestis* mtDNA [24] while the latter has only *A. artaxerxes* mtDNA. Next, we more specifically consider our study in light of the debate of what constitutes a hybrid species.

### Is *horkei* a hybrid species?

There is no consensus on what constitutes a hybrid species in the scientific literature [6,7,39– 42]. Schumer et al. [6] outlined three criteria: 1) that the hybrid species is reproductively isolated from the parental lineages, 2) that the hybrid species originated through hybridization and 3) that reproductive isolation arose as a result of hybridization. One issue with these criteria is a lack of consensus on what reproductive isolation really is, as evidenced by a recent exchange [43–46]. Another perhaps more critical problem is that a hybrid species can only arise if the parental species show incomplete reproductive isolation [39,47]. Thus, defining a hybrid species based on reproductive isolation is somewhat paradoxical [48,49]. Extremely restrictive circumstances are required for a *good* species that cannot backcross to arise from hybridization between *bad* species and persist to form its own species, especially if it has obligate sexual reproduction. To our knowledge, there is no convincing homoploid example. Instead, the reproductive isolation approach to hybrid speciation relies on partial reproductive isolation for both parental species and hybrids [40]. This is somewhat problematic since some degree of reproductive isolation is expected to be common within species [50,51], making distinction based on partial isolation arbitrary. With this in mind, we now evaluate the status of *horkei* as a hybrid species based on the three criteria outlined above. First, using population genetic data, and most decisively multispecies coalescence analysis, we have provided clear evidence for a hybrid origin of *horkei* (criterion 2). While our methods differ slightly in the inferred ancestry proportions, they all agree on i) that the minor parent ancestry is rather large and ii) that the hybridization event is neither recent nor ancient on an evolutionary timescale as inferred from e.g. the multispecies coalescence analysis and the observation of frequent ancestry switches over small scales across the genome. Thus, criterion 2 can be considered fulfilled. Second, *horkei* on Öland is geographically isolated from both parental species. The fact that *horkei* occurs in a geographic intermediate position – in which they could not have persisted since formation – suggest that this isolation should be interpreted as genetically based and therefore constitutes ecogeographic isolation. Since *horkei* is univoltine with a peak occurrence in between the two generations of *A. agestis*, they are also partially temporally isolated from *A. agestis*. On the one hand, *horkei* on the mainland (Blekingehorkei) has secondarily admixed with *A. artaxerxes* showing that these two taxa are incompletely reproductively isolated. On the other hand, *horkei* colonized Öland at least thousands of generations after the formation of the lineage without being swamped by gene flow from either parental species and has retained a sizeable portion (>30 %) of minor parental ancestry. In conclusion, *horkei* is at least partially reproductively isolated to *A. agestis* (temporal isolation) and possibly also to *A. artaxerxes*. Thus, while we have indirect evidence, direct evidence is needed before criterion 1 and by extension also criterion 3 can be considered fulfilled, but *horkei* would be considered a hybrid species under less restrictive views (e.g. patterns consistent with incomplete reproductive isolation) employed either explicitly or implicitly by several authors [7,11,13,15,16,52,53]. Genetic mapping of ecologically relevant traits (e.g. host plant preference and voltinism) would provide more information and could either support or refute the claim of *horkei* as a hybrid species. Univoltinism in *A. artaxerxes* can be broken using a long photoperiod, something that we confirmed while raising offspring for the linkage map from wild-caught females. Previous analyses of the genetic basis of voltinism variation in *Pieris napi* have identified a large-effect locus on the Z chromosome, which includes circadian clock genes known to regulate photoperiodic responses [54]. If the genetic basis of voltinism differences in *Aricia* is regulated similarly then it is possible that large effect loci reside on the Z sex chromosome which was predominantly of *A. artaxerxes* ancestry in *horkei*, in line with *horkei* being univoltine.

## 4. Conclusion

Butterflies are charismatic insects and especially the European fauna have been the subject of countless studies by scientists and naturalists for centuries. Even so, we here describe the discovery of a predominantly insular butterfly lineage that originated from a hybridization event between two non-sister species. We show that the genome of *horkei* constitutes a fine-scale mosaic of the two parental species, with varying levels of ancestry along the autosomes. However, both the mitochondrial DNA and most of the Z chromosome are derived from one of the parental species. Our results also reveal a higher level of genetic diversity in *horkei* compared to the parental species, most likely as a result of the hybridization. Finally, we show that *horkei* has persisted as a distinct lineage with a significant ancestry proportion from each parental species, despite that the original hybridization event must have occurred far from the current distribution range.

## 5. Methods

### Sampling

The sampling was planned based on information about distribution ranges from a field guide [34] and citizen science observational data from the Swedish Species Observation System (https://www.artportalen.se/). We sampled 17 *A. artaxerxes* in late June 2022 near Uppsala, central Sweden (northernmost locality in Figure 1B). We further sampled in south-eastern Sweden from 18^th^ to 25^th^ of July 2022. The latter sampling effort corresponds to an overlap between the flight periods of late flying first-generation *A. artaxerxes* and early flying second-generation *A. agestis* (Figure S1). Thirteen *horkei* individuals from Öland were sampled from three different localities. We then visited 13 locations in mainland south-eastern Sweden with previous reports of *Aricia* butterflies and found *A. artaxerxes* in one location and *A. agestis* in three locations. In addition, four *A. agestis* individuals were sampled at Tvedöra in central Skåne in late July 2022 (Table S1).

### DNA extraction and sequencing

We extracted DNA from 50 specimens using the Qiagen DNEasy Blood & Tissue kit. Samples were submitted to SciLifeLab in Stockholm, Sweden for library preparation and sequencing. Forty-five samples passed the quality control and libraries were prepared using the Illumina TruSeq PCR-free method. Samples were whole-genome sequenced on one lane of a S4 flowcell on a NovaSeq6000 instrument using 150 bp paired-end reads.

### Sequence data processing and variant calling

We filtered raw reads and removed adapter sequences using TrimGalore ver. 0.6.1, a wrapper for Cutadapt ver. 4.0 [55]. Quality of reads after filtering was assessed using MultiQC ver. 1.12 [56]. Reads were mapped to the *A. artaxerxes* reference genome (based on an individual collected in Scotland: *A. a. artaxerxes*) from the Darwin Tree of Life project, using bwa *mem* ver. 0.7.17 [57]. Mapping quality was assessed using QualiMap ver 2.2.1 [58]. We used a custom-made GATK ver. 4.3.0.0 pipeline to call variants. Variants were filtered to obtain a set of high-quality SNPs (Table S2).

### Genetic clustering analyses using SNPs

We performed two genetic clustering analyses on the high-quality SNP dataset: principal component analysis and ADMIXTURE. First, we only retained autosomal SNPs, since the genetic variation on the Z chromosome, which is present in a single copy in females, is more likely to show patterns that deviate from the general population history. Second, we filtered SNPs for pairwise linkage disequilibrium in 50 kb windows using plink ver. 2.00-alpha-3.7-20221024 [59], with an *r*^*2*^ value of allelic correlation < 0.5, obtaining 3,378,264 SNPs. We then performed a principal component analysis in plink, using the FastPCA algorithm to approximate the top 10 principal components [60]. Using the same LD-filtered dataset, we ran ADMIXTURE ver. 1.3.0 for 1 to 10 clusters (K) [61].

### Inference and analysis of Fisher junctions

It is well established that junctions between co-inherited segments (haplotype blocks) can serve as informative markers in population genetics [27,28]. Fisher junctions are the result of recombination events and are inherited and evolve much like single nucleotide polymorphisms, if a relatively recent base (or reference) population can be assumed [27]. The assumption of a relatively recent base population is required since IBD blocks will tend towards single base pairs given enough time. We developed a new method to infer Fisher junctions by using the sequenced samples as our base population. In other words, segments that are IBD were identified based on the sampled sequences and Fisher junctions were inferred in the breakpoints between IBD blocks. We inferred IBD blocks using statistical phasing of our high-quality SNP set with SHAPEIT ver. 4.2.2 [62]. For this, we assumed a constant 2 cM/Mb recombination rate along the genome. While we did indeed infer a linkage map for one of the species in this study (see below), detailed recombination rate data might be hard to obtain for many researchers and we wanted to showcase that the method is still informative when assuming a constant recombination rate. We defined a Fisher junction for a particular sample as the segment between the last variant site across the sample set and a heterozygote site with a phase different than the last heterozygote site of that particular sample (Figure S3). SHAPEIT is known to produce switch errors, i.e. erroneous junctions between haplotype blocks [26]. We therefore focused our analysis on patterns of shared Fisher junctions among samples, reasoning that switch errors will just introduce random noise and will not affect relative patterns of junction sharing. More genetically related individuals are more likely to share junctions. BEDTools ver. 2.31.1 was used to calculate overlap in Fisher junctions between samples and each overlap was counted as 1, regardless of how many base pairs that overlapped. To calculate a final overlap score (*s*) between two samples (1 and 2), we normalized the number of overlaps (*n*_*1,2*_) by the number of overlaps when each sample was compared to itself (*n*_*1,1*_ and *n*_*2,2*_):

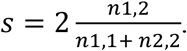

Thus, *s* ranges from 0 (no more overlap than expected randomly), to 1 (complete sharing of Fisher junctions). Since, we expected *s* to be a measure of genetic relatedness between samples, we used the KING-robust kinship estimator [63], as implemented in plink ver. 2.00-alpha-3.7-20221024 [59], for comparison and confirmed qualitative concordance between the methods. We clustered *s*-values using hierarchical clustering and visualized the patterns using the R ver. 4.3.2 [64] package corrplot [65].

### Analysis of mitochondrial DNA

We *de novo* assembled the mitochondrial DNA using GetOrganelle ver. 1.7.7.0 [66] for all individuals except two (one *A. agestis* and one *horkei* individual), for which the algorithm failed to disentangle the graph. To improve alignment efficiency, we individually annotated the mitochondrial sequences using Mitos ver. 2.1.7 [67] and rotated all sequences to begin with trnM(cat) on the + strand using Rotate ver. 1.0 [68]. We generated a multiple sequence alignment from the resulting sequences using Clustal Omega ver. 1.2 [69] enabling automatic options and trimming out small portions of repetitive content between genes and in the D-loop that poorly aligned. We generated a phylogenetic tree for the whole mitochondrial sequences using IQ-TREE ver. 2.3.0 [70] using default settings and performing 1000 ultrafast bootstraps [71] to test node support.

### Multispecies coalescence analysis

First, we randomly picked 1,000 autosomal loci, each 500 bp long, using BEDTools *random* ver. 2.29.2 [72]. We extracted haploid sequence information at each locus for each sample using the phased variants (see above). For this analysis we excluded Blekingehorkei due to indications of post-hybridization admixture from previous analyses, and thus retained 88 samples at each locus. We used the Bayesian Phylogenetics & Phylogeography (BPP) ver. 4.7.0 software to infer the evolutionary history and compare demographic scenarios under the multispecies-coalescence-with-introgression model [29]. We tested the four scenarios presented in Figure 1 of Flouri et al. [29]. In the most general model (model A), an unsampled lineage with its own coalescent events exchange alleles with the ancestor of *horkei*. This can also be seen as a complex hybrid lineage formation as opposed to the simpler hybrid lineage formation (model C). In two of the models, *horkei* branch from one of the parental species followed by unidirectional introgression from the other parental species (Model B_Age_ and B_Arx_). While all the tested multispecies coalescent models could underlie hybrid speciation events, we suggest that model C provides the strongest evidence for hybrid lineage formation since it implies that no persistent parental lineage existed prior to hybridization. For each scenario, we used a fixed species tree. Inverse gamma priors were used for the population-scaled mutation (θ) and time (τ) parameters with scale = 0.04 for θ parameters and 0.02 for τ parameters. We used a shape value of 3 (for both θ and τ), which constitutes a diffuse prior, meaning that it reduces the impact of the prior on the posterior [29]. We ran the Markov Chain Monte Carlo analysis for 1,000,000 iterations and sampled every 100 iterations, but discarded the first 50,000 iterations as burn-in. To convert coalescent time model estimates into τ (assuming one generation per year), and θ-values into population size estimates (*N*), we used the *H. melpomene* mutation rate: 2.9*10^−9^ mutations per base pair and generation [73]. A marginal likelihood (or Bayes factor) approach was used for pairwise comparisons of models [74]. To approximate the log marginal likelihood, we ran each model 16 times using the so-called power posterior, which samples from the prior to the posterior as implemented in the BFdriver function of BPP. The odds ratio of posterior probabilities of two model fits was then compared: P_1_/P_2_ = *e*^(log Likelihood Model 1 – log Likelihood Model 2)^.

### Population genetic summary statistics

We calculated population genetic summary statistics (*π, F*_*ST*_ and *D*_*XY*_) using pixy ver. 1.2.10.beta2 [75]. We used Hudson’s estimator of *F*_*ST*_ as recommended by Bhatia et al. [76]. Separate filtering was performed for the dataset used in calculating population genetic summary statistics due to the requirement of equal filtering for both variant and invariant sites for *π* and *D*_*XY*_. We excluded sites with more than 80 % missing data, and sites with lower than 10 and higher than 200 mean read depth across the entire dataset. Significant differences in π and *F*_*ST*_ between populations, as well as between genome-wide and sex chromosomes was determined by non-overlapping 95 % CI:s, a conservative measure of significance [77]. For the Z sex chromosome, only male samples (diploid for Z) were used due to constraints in how pixy handles ploidy. We tested using only the male samples for autosomes as well, but the results were only marginally affected (as expected) and therefore we used both males and females for the autosomes.

### Linkage map construction

Two of the *A. artaxerxes* individuals were caught while mating in the wild. These individuals are included in the *A. artaxerxes* group in all population genetic analyses. We raised the offspring of this cross in the laboratory using *Geranium sanguineum* and *Geranium sylvaticum* as host plants. We then extracted DNA and sequenced 40 full-sib offspring, using a bead-linked transposome complex library preparation method on Illumina NovaSeqXPlus using 150 bp paired-end reads, with technical replicates across five lanes. Reads were trimmed, mapped and deduplicated, as described above for the population resequencing data. The mean read depth across individuals ranged from 2.8x-21.8x. We inferred linkage maps for each chromosome, using Lep-MAP3 ver. 0.5 [78]. First, we used samtools *mpileup* ver. 1.19 [79] and the Pileup2Likelihood program included in Lep-MAP3 to obtain genotype likelihoods. Then we retained SNPs that 1) had no missing values, 2) did not show evidence of segregation distortion (dataTolerance = 0.001), and 3) were informative in the father (female meiosis in Lepidoptera is achiasmatic). Markers were assigned to linkage groups based on logarithm-of-the-odds likelihood value of 7 and ordered within each respective linkage group in three iterations. Despite the stringent filtering, we had thousands of markers per linkage group, which is far too many to perfectly order with the sample size of individuals in the pedigree. To solve this, we interpolated the genetic maps for each respective chromosome in R ver. 4.3.2 [64], using the R package *cobs* [80].

### Local ancestry inference

We inferred patterns of local ancestry across the genome of *horkei* using Ancestry HMM, a method that infers the local ancestry state based on a hidden Markov model, where the local ancestry is the hidden state that is inferred [31]. Since sites in physical proximity on the chromosomes are more likely to be of the same ancestry, we used the inferred linkage map to inform transition rates along the chromosomes. We also tested with twice the recombination rate inferred from the linkage map and obtained similar local allele frequencies (correlation coefficient ≈ 0.96), showing that the results are relatively robust to errors in recombination rate estimates. Only SNPs that had an allele frequency difference greater than 0.8 between *artaxerxes* and *agestis* (63, 633 SNPs on autosomes) were used as markers. We ran the analysis for the Z chromosome separately, with the same allele frequency difference, but randomly subsampled to 5,000 SNPs (to be able to run the program). We ran the Ancestry HMM analysis with the ancestry proportions (*A. artaxerxes* frequency = 0.67) estimated using ADMIXTURE as prior, using the same proportions for both the autosomes and the Z chromosome.

## Acknowledgements

The authors acknowledge support from the National Genomics Infrastructure in Genomics Production Stockholm funded by Science for Life Laboratory, the Knut and Alice Wallenberg Foundation and the Swedish Research Council, and SNIC/Uppsala Multidisciplinary Center for Advanced Computational Science for assistance with massively parallel sequencing and access to the UPPMAX computational infrastructure. J.B acknowledges funding from Lennanders stiftelse and the Birgitta Sintring foundation. N.B acknowledges funding from the Swedish Research Council (project grant 2019-04791). We thank Mahwash Jamy and Melina Eberhagen for help with sampling. We also thank Alexander Mackintosh, Axel Jensen and Zachariah Gompert for insightful discussions.

## Data availability statement

Whole-genome sequencing data generated for this study is available on ENA under accession: PRJEB81148. Analysis scripts is available in the following repository on GitHub: https://github.com/JesperBoman/Horkei_begins.

## Author contributions

JB, ZN and NB designed research. JB, ZN and NB performed research. JB and ZN analysed data and JB wrote the paper with feedback from ZN and NB.

