## Supplementary Information for "On the origin of an insular hybrid butterfly lineage"

**Figure S1:** Distribution of observations of *A. artaxerxes* and *A. agestis* in Sweden from 2000-01-01 – 2023-11-01 downloaded from Artportalen, the Swedish species observation system (<https://www.artportalen.se/>). *A. artaxerxes* and *A. agestis* are partially temporally isolated with the two generations of *A. agestis* occurring before and after the flight peak of *A. artaxerxes.*

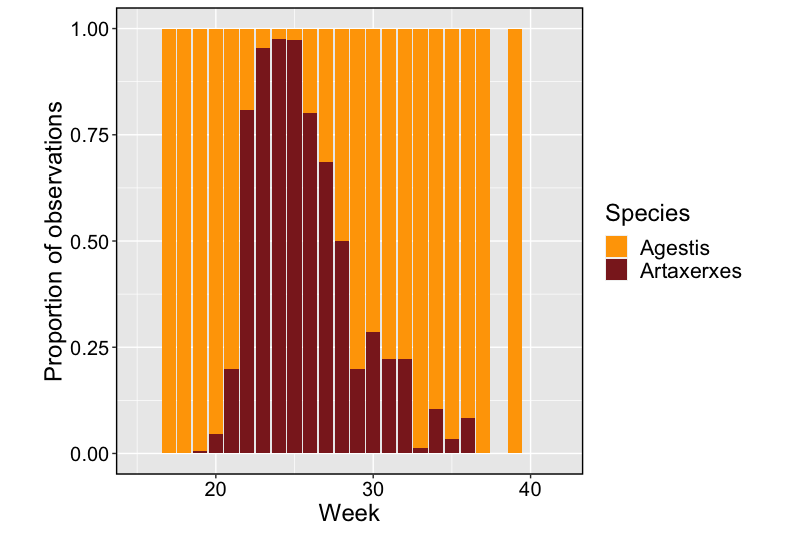

**Figure S2:** Cross-validation error analysis based on the ADMIXTURE results. The cross-validation error shows a monotonic increase with the lowest value for K = 1. This highlights the large proportion of shared variation in this group of butterflies.

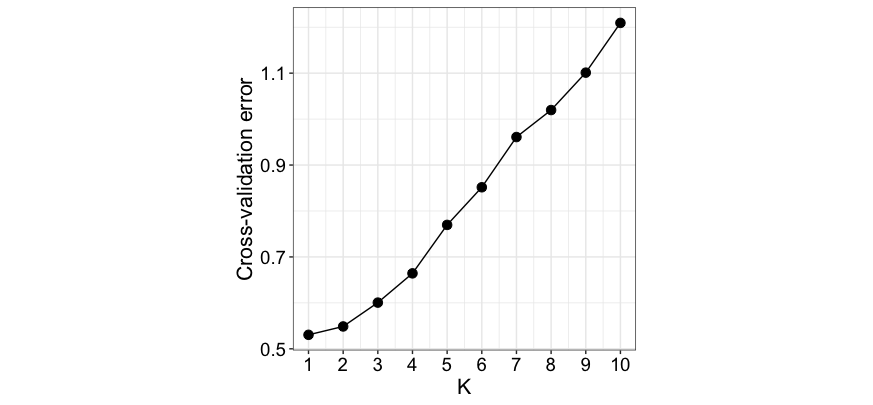

**Figure S3:** From phased genotypes to Fisher junctions. **(A)** Representation of IBD segments (haplotype blocks) inferred using statistical phasing of biallelic SNPs in a selected region of chromosome 1. 0|0 and 1|1 represent genotypes homozygous for the reference allele and the alternative allele, respectively. For readability, the plot only shows variant sites across the whole sample set. In this region, there was little variation in the *A. artaxerxes* population (top section). **(B)** The regional lack of heterozygosity in the *A.artaxerxes* population is reflected by long haplotypes and few Fisher junctions, compared to the other sample groups.

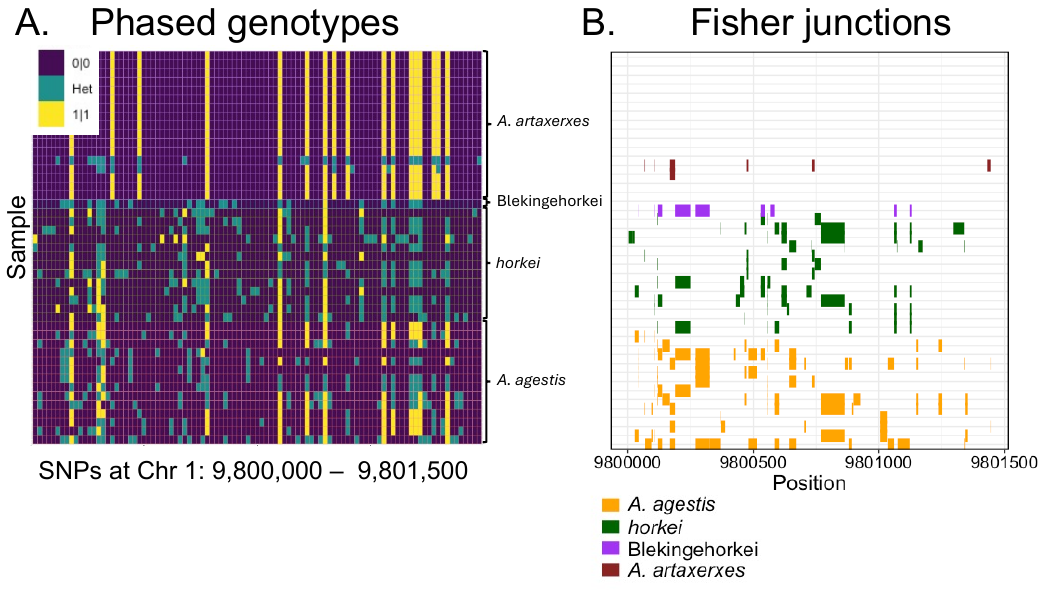

**Figure S4:** Genetic relatedness among samples. We used the KING-robust kinship estimator to estimate genetic relatedness between samples**.** Some relationships like clustering of distant relatives at the KBH locality (column names) are similar to the Fisher junction analysis. However, the KING-robust kinship estimator does not capture the weaker effects of fine-scale structure among *agestis*-localities.

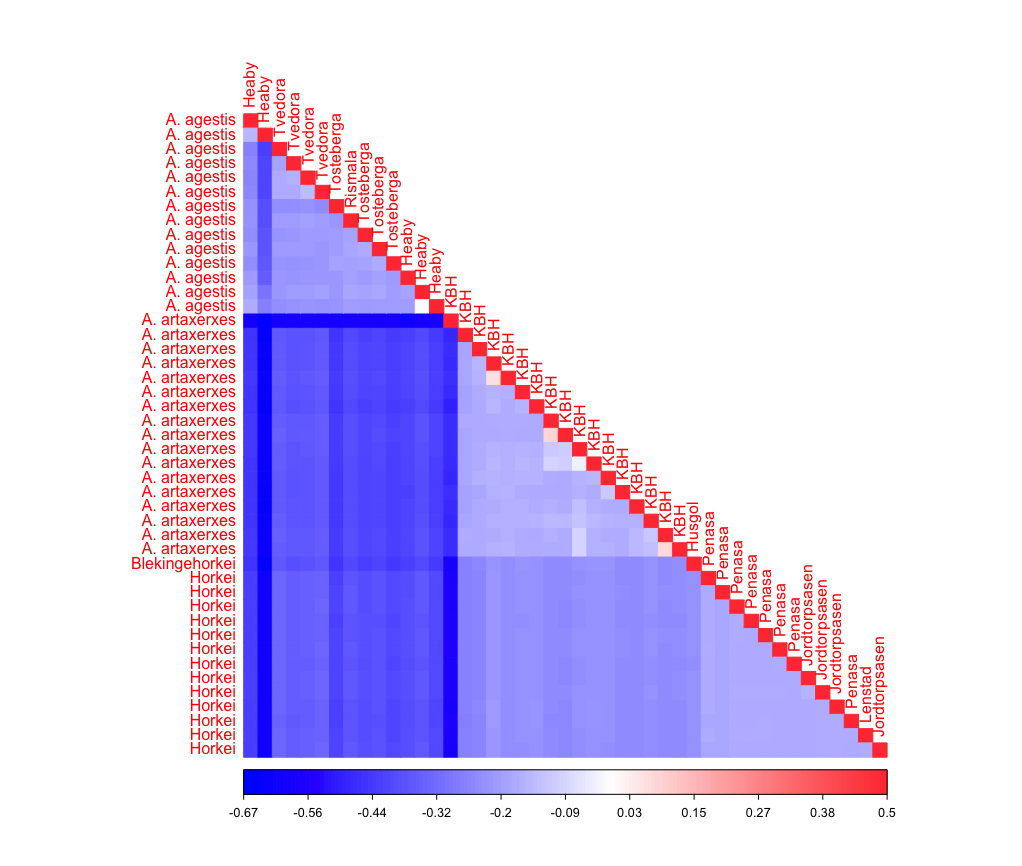

**Figure S5:** Venn diagram of *agestis* alleles segregating in admixed population. We inferred autosomal fixed differences between *A. artaxerxes* and *A. agestis* (n = 5,759). We then overlapped *A. agestis* alleles present in *horkei* and Blekingehorkei. Only 7 *A. agestis* alleles were present in Blekingehorkei but absent in *horkei.* This small number could easily be a result of genetic drift or sampling effects and thus indicates that *agestis* alleles in these two admixed populations are derived from the same hybridization event.

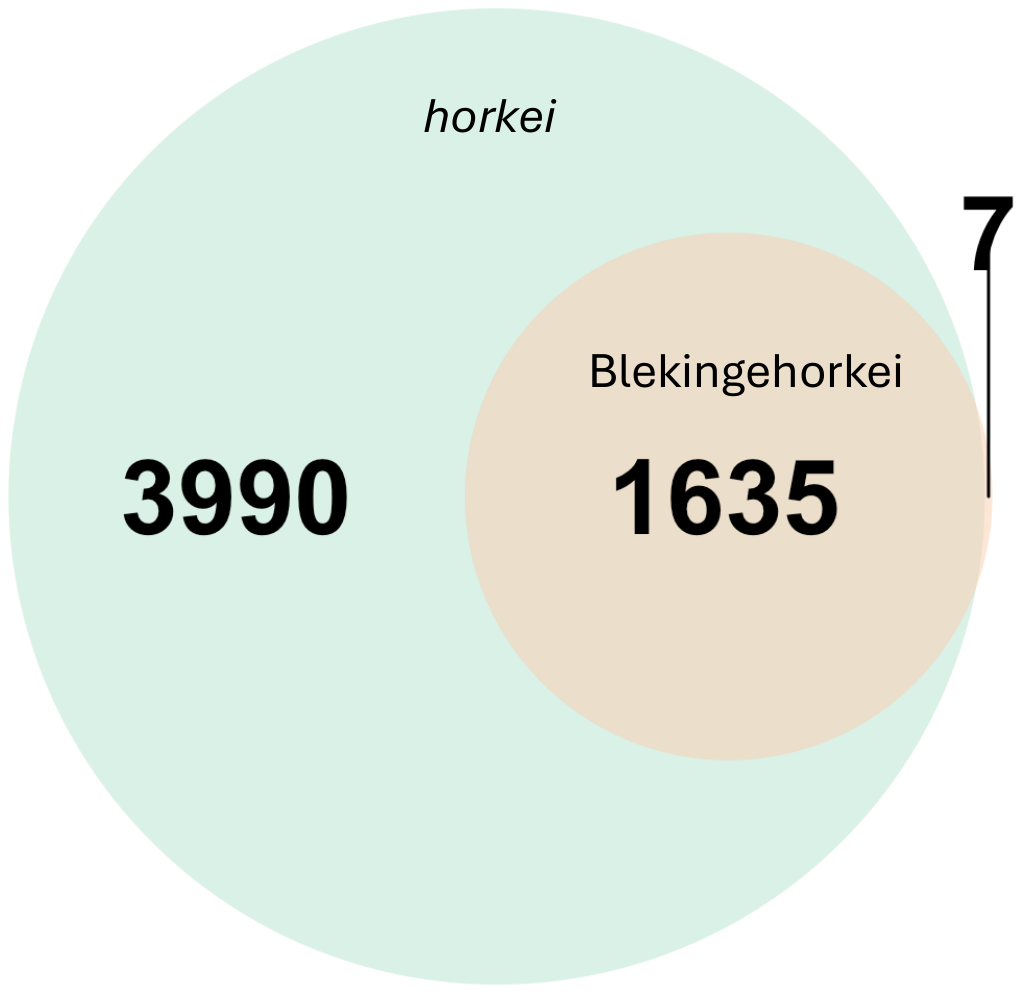

**Figure S6:** Genetic differentiation (*F_ST_*) between *A. agestis* and *A. artaxerxes.* Chromosome 23 is the Z sex chromosome.
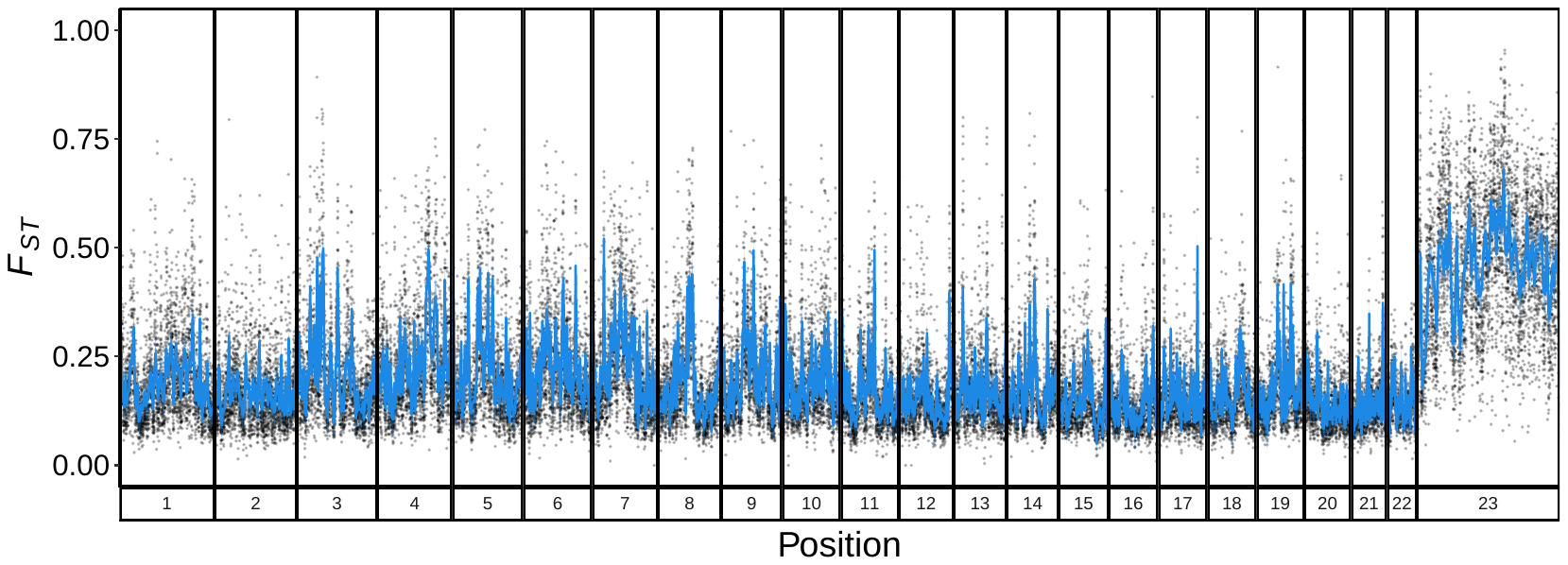

**Figure S7:** Marey map of each linkage group. We used a cohort of full-sib offspring to generate a linkage map. Average male map length per chromosome was ~51 cM, or one expected crossover per chromosome per male meiosis. Y-axis shows the male genetic map length, and the X-axis shows the physical map length. Chromosome 23 is the Z sex chromosome. Black points are the inferred map position, while red lines trace the interpolated map used for local ancestry analysis. Interpolation was used to reduce influence of a few disordered markers on the final genetic map. These markers are most likely disordered due to the large total number of markers used (as a result of whole-genome resequencing).

**Table S1.** List of samples. ID stands for identification. The number of reads is given in millions. Sex was determined by coverage analysis of the Z sex chromosome, in which females have reduced coverage of the ancestral Z region compared to males.

| **ID** | **Reads** | **Locality** | **Region** | **Taxon** | **Sex (Cov.)** |
| --- | --- | --- | --- | --- | --- |
| 184 | 44.72 | Heaby | Blekinge | *A. agestis* | F |
| 111 | 80.50 | Heaby | Blekinge | *A. agestis* | M |
| 113 | 62.28 | Heaby | Blekinge | *A. agestis* | M |
| 136 | 63.73 | Heaby | Blekinge | *A. agestis* | M |
| 160 | 49.64 | Heaby | Blekinge | *A. agestis* | M |
| 112 | 53.39 | Rismala | Smaland | *A. agestis* | M |
| 144 | 53.13 | Tosteberga | Skane | *A. agestis* | M |
| 152 | 52.70 | Tosteberga | Skane | *A. agestis* | M |
| 168 | 47.52 | Tosteberga | Skane | *A. agestis* | M |
| 176 | 49.69 | Tosteberga | Skane | *A. agestis* | M |
| 142 | 66.98 | Tvedora | Skane | *A. agestis* | M |
| 150 | 54.89 | Tvedora | Skane | *A. agestis* | M |
| 158 | 62.15 | Tvedora | Skane | *A. agestis* | M |
| 166 | 51.75 | Tvedora | Skane | *A. agestis* | M |
| 102 | 64.66 | KBH | Uppland | *A. artaxerxes* | F |
| 118 | 61.93 | KBH | Uppland | *A. artaxerxes* | F |
| 125 | 51.80 | KBH | Uppland | *A. artaxerxes* | F |
| 126 | 119.76 | KBH | Uppland | *A. artaxerxes* | F |
| 133 | 53.44 | KBH | Uppland | *A. artaxerxes* | F |
| 134 | 55.83 | KBH | Uppland | *A. artaxerxes* | F |
| 141 | 77.31 | KBH | Uppland | *A. artaxerxes* | F |
| 101 | 58.60 | KBH | Uppland | *A. artaxerxes* | M |
| 109 | 50.39 | KBH | Uppland | *A. artaxerxes* | M |
| 110 | 62.20 | KBH | Uppland | *A. artaxerxes* | M |
| 117 | 68.25 | KBH | Uppland | *A. artaxerxes* | M |
| 149 | 52.30 | KBH | Uppland | *A. artaxerxes* | M |
| 157 | 54.04 | KBH | Uppland | *A. artaxerxes* | M |
| 165 | 50.51 | KBH | Uppland | *A. artaxerxes* | M |
| 173 | 54.62 | KBH | Uppland | *A. artaxerxes* | M |
| 181 | 48.32 | KBH | Uppland | *A. artaxerxes* | M |
| 189 | 52.81 | KBH | Uppland | *A. artaxerxes* | M |
| 192 | 88.93 | Husgol | Blekinge | *Blekingehorkei* | F |
| 120 | 59.23 | Jordtorpsasen | Öland | *Horkei* | F |
| 119 | 59.21 | Jordtorpsasen | Öland | *Horkei* | M |
| 128 | 58.52 | Jordtorpsasen | Öland | *Horkei* | M |
| 174 | 63.17 | Jordtorpsasen | Öland | *Horkei* | M |
| 127 | 65.74 | Lenstad | Öland | *Horkei* | M |
| 151 | 54.87 | Penasa | Öland | *Horkei* | F |
| 105 | 53.71 | Penasa | Öland | *Horkei* | M |
| 135 | 54.39 | Penasa | Öland | *Horkei* | M |
| 159 | 59.72 | Penasa | Öland | *Horkei* | M |
| 167 | 55.34 | Penasa | Öland | *Horkei* | M |
| 175 | 62.13 | Penasa | Öland | *Horkei* | M |
| 183 | 55.47 | Penasa | Öland | *Horkei* | M |
| 191 | 48.03 | Penasa | Öland | *Horkei* | M |

**Table S2.** Filtering parameters for population-resequencing data to obtain a set of high-quality SNPs.

| **Parameter** | **Filtering threshold** |
| --- | --- |
| Fisher strand bias | <60 |
| Strand odds ratio | <3 |
| Mapping quality | >40 |
| Mapping quality rank sum test | >-12.5 |
| Quality by depth | >2 |
| Read position rank sum test | >-8 |
| Depth | Less than 3 standard deviations above mean read coverage per variant site |
| Max missing | 1 (no missing allele information) |
